# Complementary cytotoxicity of GD2-targeted photoimmunotherapy and 5-aminolevulinic acid photodynamic therapy in neuroblastoma and osteosarcoma

**DOI:** 10.64898/2026.09.01.748695

**Authors:** Arjun Pant, Catherine Yip, Bhuvitha Chagantipati, Milena Mattioli, Virginia M. Kelly, Eesa Noaman, Arianna Kahler-Quesada, Brady Grano-Mickelsen, Sydney A. Jackson, Peter L. Choyke, Hisataka Kobayashi, Costas G. Hadjipanayis, Lauren T. Rosenblum, Marcus M. Malek, Gary Kohanbash

**Affiliations:** Department of Surgery, University of Pittsburgh, Pittsburgh, PA; Department of Neurological Surgery, University of Pittsburgh, Pittsburgh, PA; Molecular Imaging Program, National Cancer Institute, NIH, Bethesda, MD, USA

## Abstract

Phototherapy, a light-activated anticancer treatment, enables localized tumor-cell killing with distinct mechanisms of action. Photoimmunotherapy (PIT) produces immunogenic tumor cell death upon near-infrared light activation of a photoabsorber through antigen-specific targeting. Photodynamic therapy (PDT) produces reactive oxygen species through red-light activation of intracellular protoporphyrin IX generated from 5-aminolevulinic acid uptake and metabolism. PIT may have limited activity in antigen-low cells, whereas PDT has less precise tumor selectivity. We combined these modalities to define their interaction, broaden cytotoxicity, and determine whether dual treatment could reduce light-dose requirements. We conjugated dinutuximab, which targets the GD2 antigen, to IRDye 700DX and characterized plasma-membrane localization by confocal and widefield microscopy. PIT and PDT monotherapies were evaluated across agent and light doses in neuroblastoma (NB) and osteosarcoma (OS) cell lines. Combination matrices were tested using interaction, highest-single-agent, and Bliss analyses. Both monotherapies demonstrated significant light-dose-dependent effects in NB and OS. PIT produced no measurable cytotoxicity in antigen-blunted control cells, whereas PDT remained effective, confirming antigen-dependence of PIT and antigen-independence of PDT. The combination interaction was significant in SK-N-BE(2) but not LM7. At selected combinations, however, dual treatment produced greater killing than the more effective matched monotherapy in both SK-N-BE(2) and LM7 (P_adj_≤0.022). Notably, lowest combination of PIT 10 J/cm² plus PDT 10 J/cm^2^ achieved 90.3% killing in SK-N-BE(2), exceeding higher light-dose PIT or PDT monotherapy, suggesting a light-dose sparing effect. These findings establish potent and complementary PIT-PDT activity, supporting dual phototherapy to broaden cytotoxicity and reduce light-dose requirements in GD2-expressing tumor phototherapy.

## INTRODUCTION

Neuroblastoma (NB) and osteosarcoma (OS) are among the most clinically challenging pediatric extracranial solid tumors. Among these, NB is the most common and disproportionately accounts for approximately 15% of pediatric cancer deaths (1). Surgical resection is an important component of local control. However, primary tumors frequently encase major vessels or involve anatomically complex regions which limits the ability to perform safe and complete resection, potentially resulting in substantive operative morbidity (2–4). Small, occult tumors may also evade intraoperative detection, contributing to residual disease (2). Surgeon-assessed resection is less than 90% in approximately 30% of patients (5). Patients with high-risk disease therefore receive intensive multimodal therapy consisting of chemotherapy, radiotherapy, and immunotherapy (1, 6, 7). Despite this, relapse remains frequent, with poor outcomes. In a large international cohort of patients with first relapse or progression, 5-year overall survival was only 20% (8).

OS is similarly challenging to treat. Surgery is the cornerstone of therapy, yet complete resection can be challenging, and outcomes for metastatic, recurrent, or unresectable disease have not improved substantially despite multiagent chemotherapy (9). Metastatic disease also portends a poor prognosis with a median survival of 8.4 months and virtually no survival for patients whose lungs cannot be cleared of disease (10). In a Children’s Oncology Group analysis, 5-year survival after first recurrence was 17.7% (11). An alternative, locally activated therapy could therefore address two important limitations of current therapy: eliminating residual or surgically inaccessible tumor and serving as a therapeutic adjunct for tumors resistant to or less susceptible to other treatment modalities.

Near-infrared photoimmunotherapy (PIT) is a selective, molecularly targeted anticancer therapy that couples a tumor-directed antibody to the near-infrared photoabsorber IRDye 700DX (IR700). After membrane binding of the antibody-photoabsorber conjugate (APC), near-infrared illumination releases the hydrophilic axial ligands of IR700, producing a marked loss of hydrophilicity for the entire molecule. The resulting APC complexes aggregate, producing focal plasma-membrane disruption (Fig. 1), osmotic swelling, and necrotic cell death (12–15).

**Figure 1.**
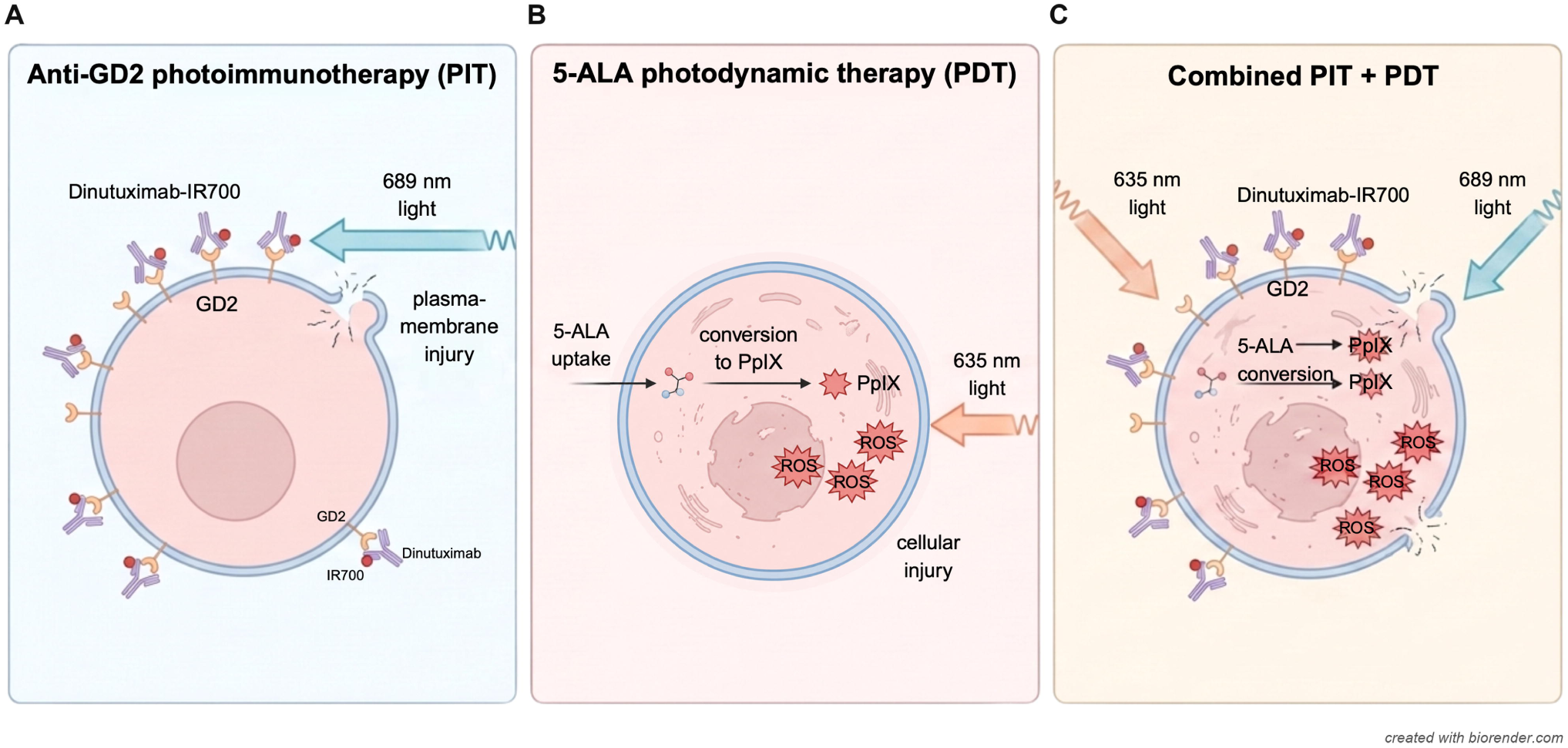
Mechanisms of PIT, PDT and combination phototherapy. (A) Dinutuximab-IR700 binds cell-surface GD2 and, after 689-nm illumination, produces plasma-membrane injury. (B) Cellular uptake of 5-aminolevulinic acid (5-ALA) and conversion to intracellular protoporphyrin IX (PpIX), followed by 635-nm illumination, generates ROS and produces cellular injury. (C) Combined PIT and PDT activates membrane-bound dinutuximab-IR700 and intracellular PpIX using their respective excitation wavelengths to produce plasma-membrane injury and ROS production.

Cytotoxicity therefore requires both target binding and illumination. Clinical feasibility of PIT was demonstrated in a phase 1/2a study of cetuximab-IR700 targeting epidermal growth factor receptor, and the conjugate was subsequently approved in Japan for unresectable locally advanced or recurrent head and neck cancers (16, 17).

Disialoganglioside GD2 is highly expressed on most NB tumors and on a vast majority of OS tumors (18, 19). Expression in normal tissue is substantially lower and is concentrated principally in peripheral sensory nerve fibers, melanocytes and other neural tissues (19). Given the selective tumor expression of GD2, PIT using monoclonal anti-GD2 APCs has demonstrated promising preclinical efficacy against both NB and OS (20–22).

Dinutuximab is a U.S. Food and Drug Administration (FDA)-approved anti-GD2 monoclonal antibody and standard-of-care maintenance immunotherapy for children with high-risk NB who respond to initial multimodal therapy (6, 7). Conjugation of dinutuximab to IR700 may therefore enable targeted GD2 binding and cytotoxic activity restricted to the illuminated field. This approach uses a clinically established, FDA-approved antibody backbone and provides a translationally relevant starting point.

While the molecularly targeted nature of PIT is its principal strength, it also carries the potential limitation of reduced efficacy in tumors with heterogeneous antigen expression. An alternative phototherapy, photodynamic therapy (PDT) using 5-aminolevulinic acid (5-ALA), is an established anticancer treatment. It provides a complementary, antigen-independent phototherapy mechanism: exogenous 5-ALA enters the heme biosynthetic pathway and promotes intracellular accumulation of protoporphyrin IX (PpIX), which generates reactive oxygen species (ROS) after illumination (23). Unlike antibody-directed PIT, 5-ALA PDT does not require a specific surface antigen and has been investigated across multiple oncologic applications (24). Preclinical studies have also demonstrated 5-ALA-mediated phototoxicity in NB and bone sarcoma cell lines (25–27). However, 5-ALA/PpIX is only partially selective as non-neoplastic and inflammatory tissues can absorb 5-ALA and exhibit PpIX fluorescence (24). Despite this, the 10-patient phase I INDYGO study supports long-term safety after intraoperative 5-ALA PDT of the glioblastoma resection cavity, with no treatment-related deaths or brain necrosis on follow-up (28, 29). Normal tissue injury is also a consideration for PIT, as excessive near-infrared illumination can cause thermal injury, edema and bleeding (30).

Combining PIT with PDT presents a unique opportunity to broaden cytotoxicity in antigen- and treatment-heterogeneous tumors that are less sensitive to either modality alone. Their complementary effects may further reduce the illumination required for phototoxicity.

In this study, we characterized and evaluated two phototherapies for NB and OS: GD2-directed PIT based on a dinutuximab backbone and established, antigen-independent 5-ALA PDT. We examined treatment effects across a range of agent and light doses, assessed antigen dependence, and identified factors influencing the magnitude and kinetics of treatment response. Finally, we evaluated these mechanistically distinct therapies in combination, focusing on their interaction, implications for reduced light dosing, and improved overall extent of cytotoxicity.

## MATERIALS AND METHODS

### *In vitro* maintenance of cell lines

Human NB SK-N-BE(2) cells (ATCC; RRID: CVCl_0528) and NMB6 cells (Ira Bergman laboratory, University of Pittsburgh; derived from NMB RRID: CVCL_2143) were cultured in minimum essential medium alpha (MEM-alpha; Gibco, cat. #12-571-048) supplemented with 10% heat-inactivated fetal bovine serum (Gemini Bio, cat. #100-106-500), Normocin (InvivoGen, cat. #ant-nr-2), antibiotic-antimycotic (Gibco, cat. #15-240-062), L-glutamine (Gibco, cat. #25-030-081), and MEM nonessential amino acids (Gibco, cat. #11140-050). Human OS 143B (ATCC; RRID: CVCL_2270) and LM7 cells (Kurt Weiss laboratory, University of Pittsburgh; RRID: CVCL_0515) were cultured in Dulbecco’s modified Eagle medium (DMEM) without glutamine (Corning, cat. #MT10017CV) supplemented with 10% heat-inactivated fetal bovine serum, penicillin-streptomycin (Gibco, cat. #15-140-122), MEM vitamin solution (Gibco, cat. #11120052), and MEM nonessential amino acids (Gibco, cat. #11-140-050). SF8628 (University of California, San Francisco Medical Center; RRID: CVCL_IT46) diffuse midline glioma and U-87 MG (ATCC; RRID: CVCL_0022) glioblastoma (GBM) were included in a GD2-expression study, and HEK293T (ATCC; RRID: CVCL_0063) cells were used as a GD2-negative PIT and PDT comparator. SK-N-BE(2) and 143B derivatives with incidentally blunted GD2 expression were generated in-house by transfection with the firefly luciferase gene. All cell lines were human-derived, obtained in 2020 from the sources listed above, and used within 10 passages from thawing. SK-N-BE(2), 143B, U-87 and HEK293T cell lines were authenticated through genotypic analysis performed by ATCC. All cell lines tested negative for mycoplasma prior to first use. SK-N-BE and U87 were derived from male donors. NMB6, LM7, 143B, SF8628 and HEK293T were derived from female donors. Cell lines were maintained at 37°C in a humidified incubator containing 5% CO_2_.

### Dinutuximab-IRDye tracer conjugation and evaluation

Dinutuximab was conjugated to IR700 or IRDye 800CW (IR800) using an amine-reactive N-hydroxysuccinimide (NHS) ester modified from a method previously described by our lab (2). IRDye 700DX NHS Ester was provided by the Kobayashi laboratory (National Institutes of Health, Bethesda, MD; previously sold by LI-COR Biosciences, cat. #929-70010), and IRDye 800CW NHS Ester was obtained from LI-COR Biosciences, Lincoln, NE (cat. #929-70020).

Briefly, dinutuximab (Unituxin; United Therapeutics Corporation, Research Triangle Park, NC; RRID: AB_3695149; 1 mg, 6.7 nmol) was incubated with a fivefold molar excess of IRDye 700DX NHS Ester or a threefold molar excess of IRDye 800CW NHS Ester in 0.1 mol/L sodium carbonate buffer at pH 8.5-9.0 for 3 hours at room temperature under light-protected conditions. Unconjugated photoabsorber was removed by centrifugal ultrafiltration using a 30-kDa molecular-weight-cutoff concentrator (Pierce Protein Concentrator; Thermo Fisher Scientific, cat. #88522). Protein and photoabsorber absorbance were measured at 280 nm and at the corresponding photoabsorber absorbance maxima (689 nm for IR700 and 777 nm for IR800) by ultraviolet-visible (UV-Vis) spectrophotometry (NanoDrop One, Thermo Scientific; RRID: SCR_023005) to calculate the number of IR700 or IR800 per antibody. Conjugates were stored in Dulbecco’s phosphate-buffered saline (PBS; Corning, cat. #21-031-CV) at 1.8 mg/mL at 4 °C in the dark. Selected IR700 and IR800 conjugate preparations were analyzed by size-exclusion high-performance liquid chromatography (SEC-HPLC) on an Agilent Technologies 1260 Infinity (RRID: SCR_019360) system with absorbance monitoring at 254 and 280 nm.

### Flow-cytometric quantification of GD2 expression

GD2 expression was measured using a BD LSRFortessa flow cytometer (BD Biosciences, RRID: SCR_018655) and BD FACSDiva software (RRID: SCR_001456). A total of 200,000 cells from each NB, OS, and glioma cell line were washed in cold staining buffer consisting of PBS with 1% bovine serum albumin (Fisher Scientific, cat. # BP1605-100) before brief incubation with 1 µL Human TruStain FcX (BioLegend, cat. #422302) to prevent nonspecific antibody binding. Cells were subsequently stained with 1 µg phycoerythrin (PE)-conjugated anti-GD2 antibody (clone 14G2a; BioLegend, cat. #357303) for 1 hour at 4 °C. A standard curve of fluorescence was generated using BD QuantiBRITE beads with known levels of PE staining, as described in the PE fluorescence quantitation kit (BD Biosciences, cat. #340495).

Data were analyzed in FlowJo version 10 (RRID: SCR_008520). The estimated GD2 expression per cell was calculated by substituting geometric mean PE-area fluorescence intensity of GD2-positive singlets into the logarithmic standard curve equation.

### Photoimmunotherapy and photodynamic therapy

Cells were seeded in 24-well plates at 75,000 cells per well with 1 mL of corresponding culture medium. Cells were incubated with 5-ALA (MedChemExpress, USA, cat. #5451-09-2) for 48 hours and/or dinutuximab-IR700 for 5 hours at 37°C, depending on the allocated treatment. Prior to illumination, culture medium was replaced with 1 mL of the corresponding phenol red-free complete growth medium.

Illumination was delivered with a phototherapy platform (Modulight, Tampere, Finland, model ML8500) in well-by-well mode at an irradiance of 150 mW/cm². Dinutuximab-IR700 photoimmunotherapy (PIT) was performed at 689 nm, and 5-ALA photodynamic therapy (PDT) was performed at 635 nm. Negative controls included untreated cells, light-only controls where applicable, and agent-treated wells that did not receive illumination (0 J/cm²). Hydrogen peroxide (CVS Health, item #209478; 1500µM, 24 hour incubation) and camptothecin (Thermofisher Scientific, cat. #J62523.MD; 72 µM, 24 hour incubation) were included as necrosis and apoptosis positive controls, respectively. All groups were treated and analyzed in replicates of three or more.

For monotherapy PIT light-dose experiments, cells were incubated with a fixed 15 µg/mL dinutuximab-IR700 agent dose and exposed to 0, 10, 20, 50, 75, or 100 J/cm² light at 689 nm. For PIT agent-dose experiments, NMB6 and SK-N-BE(2) cells received 0, 5, 10, 15, or 20 µg/mL dinutuximab-IR700 at a fixed light dose of 50 J/cm²; LM7 was tested at 0, 5, 10, 15, 20, and 30 µg/mL. For PDT light-dose experiments, cells were incubated with a fixed 500 µmol/L 5-ALA and exposed to 0, 10, 30, 50, or 75 J/cm² at 635 nm. For PDT agent-dose experiments, cells received 0, 500, 1,000, 1,500, or 2,000 µmol/L 5-ALA and either 0 or 30 J/cm². The monotherapy experimental conditions are summarized in Supplementary Table S1.

Combination experiments used fixed agent doses of 10 µg/mL dinutuximab-IR700 and 500 µmol/L 5-ALA. PIT illumination was performed at 0, 10, 20, or 50 J/cm², and PDT at 0, 10, or 30 J/cm². The 689-and 635-nm sources were programmed within the same ML8500 well protocol and were treated as concurrent exposures. PIT and PDT monotherapy were performed within the same experiment for comparison, and these wells received only the wavelength corresponding to that treatment.

For the HEK293T comparator non-cancer control experiment, cells received 10 µg/mL dinutuximab-IR700 with 20 J/cm² at 689 nm or 500 µmol/L 5-ALA with 10 J/cm² at 635 nm. Comparator experiments with dinutuximab-IR800 used 810-nm illumination at 20, 50, or 100 J/cm². Untreated wells served as the same-run control for both comparator experiments.

A time-course experiment to assess the kinetics of cell death post PIT or PDT was performed with SK-N-BE(2) cells using 10 µg/mL dinutuximab-IR700 and 20 J/cm² light or 500 µmol/L 5-ALA and 10 J/cm² light. Cells were collected 1, 24, or 48 hours after illumination. Cell viabilities were normalized to the mean viability of the control wells collected at the corresponding time point.

Mechanistic experiments examining culture-medium dependence were performed in SK-N-BE(2) cells using same agent concentrations and light doses as in the time-course experiment. For the no-medium conditions, culture medium was aspirated immediately before PIT/PDT (cells were treated in the complete absence of any medium) and restored immediately afterward. The corresponding medium-present conditions were illuminated with no growth medium removal.

### Flow-cytometric assessment of cell viability and death

Cell viability was assessed at 24 hours across all experiments except the time-course experiment, for which the collection times are described above. Flow cytometry with allophycocyanin (APC)-conjugated Annexin V and propidium iodide (PI) stains (BioLegend, cat. #640932) was used to quantify cell death and characterize the type of death. Culture supernatants were retained to include detached cells. Adherent cells were detached with 0.05% trypsin (Corning, cat. #MT25052CI) and pooled with the corresponding supernatant. Cells were washed with Annexin V binding buffer (cat. #422201) and resuspended in 100 µL binding buffer containing 5 µL APC Annexin V. After a 15-minute incubation at room temperature in the dark, cells were washed and resuspended in binding buffer. PI (2 µL) was added immediately before acquisition on the BD LSRFortessa flow cytometer.

Unstained, Annexin V-only, and PI-only controls were used to establish gates and confirm channel separation. Data were analyzed on FlowJo version 10. Annexin V-negative/PI-negative events were classified as viable cells. Annexin V-positive, PI-positive, and double-positive events were classified as nonviable cells.

The primary endpoint measured was the viable fraction, defined as the percentage of Annexin V-negative and PI-negative flow-cytometry events for each group. Treatment response was reported as viability relative to the mean viability of the corresponding same-run negative-control wells. Each experimental condition was tested in at least three replicates using separately plated, treated, and processed wells, across all experiments.

### Microscopy

Widefield IR700 fluorescence images were acquired through the Cy5 channel on an ECHO Revolve microscope (RRID: SCR_026523) with a 20x objective. Identical acquisition settings were used for all conditions within each comparison. At least five randomly selected clusters containing cells were evaluated per condition across images. Total cluster fluorescence (TCF) was corrected for background fluorescence. Mean fluorescence intensity (MFI) was further quantified as TCF per unit area of cluster using ImageJ software (RRID: SCR_003070). SK-N-BE(2), LM7, 143B and U87 were imaged post-incubation with 15 µg/mL dinutuximab-IR700 for 5h. Additional fluorescence experiments with SK-N-BE(2) cells assessed signal before and at 1 and 24 hours after PIT, imaging cells with and without culture medium, 2 IR700 per antibody versus 3, and 5 versus 20 µg dinutuximab-IR700.

Confocal z-stacks were acquired on a Nikon A1R confocal (RRID:SCR_020317) with total internal reflection fluorescence system with a 100× objective. SK-N-BE(2) and 143B cells were stained for 1 hour with CellMask Orange Plasma Membrane Stain (Thermo Fisher Scientific, cat. #C10045) after incubation with dinutuximab-IR700 for 5 hours. Dinutuximab-IR700 and CellMask signals were acquired in the 640- and 561-nm channels, respectively.

Brightfield images were acquired with a 10x objective on an EVOS FL Auto 2 microscope (Thermo Fisher Scientific, cat. #AMAFD2000). The main brightfield comparisons were obtained 24 hours after treatment.

### Statistical analysis

Treatment levels were modeled as categorical variables. Single-agent experiments were evaluated using ordinary least-squares (OLS) models with HC3 heteroskedasticity-consistent covariance estimates. Within each cell line, an omnibus Wald test assessed differences across all tested treatment levels. Holm adjustment was applied across cell-line omnibus tests (four for the light-dose experiments and three for the PIT agent-dose experiments) and to prespecified dose versus reference contrasts within each cell line. Reported P values describe separation among wells within the corresponding experimental run.

The primary combination-matrix analysis used a factorial model containing categorical PIT/PDT illuminations, and their interaction among dual-agent wells. Viable proportions were modeled on the logit scale with HC3 covariance estimates. Untransformed viable percentages were examined in a sensitivity analysis. The two cell-line interaction tests were Holm adjusted.

Combination performance was also evaluated relative to same run monotherapies using highest-single-agent (HSA) and Bliss models. HSA killing was calculated as the viability of the more effective pure monotherapy minus combination viability, reported as control normalized percentage points. Bliss independence used the same-run untreated control (V00), pure PIT monotherapy at PIT light dose i (Vi0), and pure PDT monotherapy at PDT light dose j (V0j).

Expected raw viability for the combination at i,j was (Vi0 × V0j)/V00. Bliss excess killing was calculated as expected minus observed viability, normalized to V00, and reported in percentage points. Bliss killing was interpreted descriptively: negative values indicated less killing than expected, and positive values indicated greater killing (synergy). Values close to zero were considered consistent with Bliss additivity.

For comparisons between parental and GD2-blunted derivatives, calibrated PE estimates were log2 transformed, and exponentiated model coefficients provided fold differences between parent and GD2-blunted cell lines. Comparisons used HC3 covariance estimates with Holm adjustment. The relationship between GD2 abundance and PIT killing at 50 J/cm² across the four parental cell lines was summarized using Spearman rank correlation coefficient (ρ). IR700 MFI per unit area was compared among NB, pooled OS (LM7 and 143B), and GBM image regions using Welch’s one-way analysis of variance (ANOVA).

Medium-condition and time-course experiments were analyzed using HC3 factorial models containing treatment modality, experimental condition or time, and their interaction. Time course wells were normalized to corresponding time-matched untreated control. Prespecified within-modality contrasts were adjusted using the Holm method. Tests across all experiments were two-sided with alpha = 0.05 and were performed in R version 4.5.2.

### Data availability

The data generated in this study are available from the corresponding author on reasonable request.

## RESULTS

### Chemical and fluorescence properties of antibody-photoabsorber conjugate

SEC-HPLC chromatography of dinutuximab-IR700 and dinutuximab-IR800 was performed to assess size distribution, aggregation and impurities in the conjugation product. Each demonstrated a dominant conjugate-associated peak at approximately 6 minutes and a comparatively smaller impurity-associated peak near 9 minutes (Fig. 2A and B). Using UV-vis spectrophotometry, the number of IR700 and IR800 per antibody were calculated to be 1.94 ± 0.469 and 2.20 ± 0.277, respectively.

**Figure 2.**
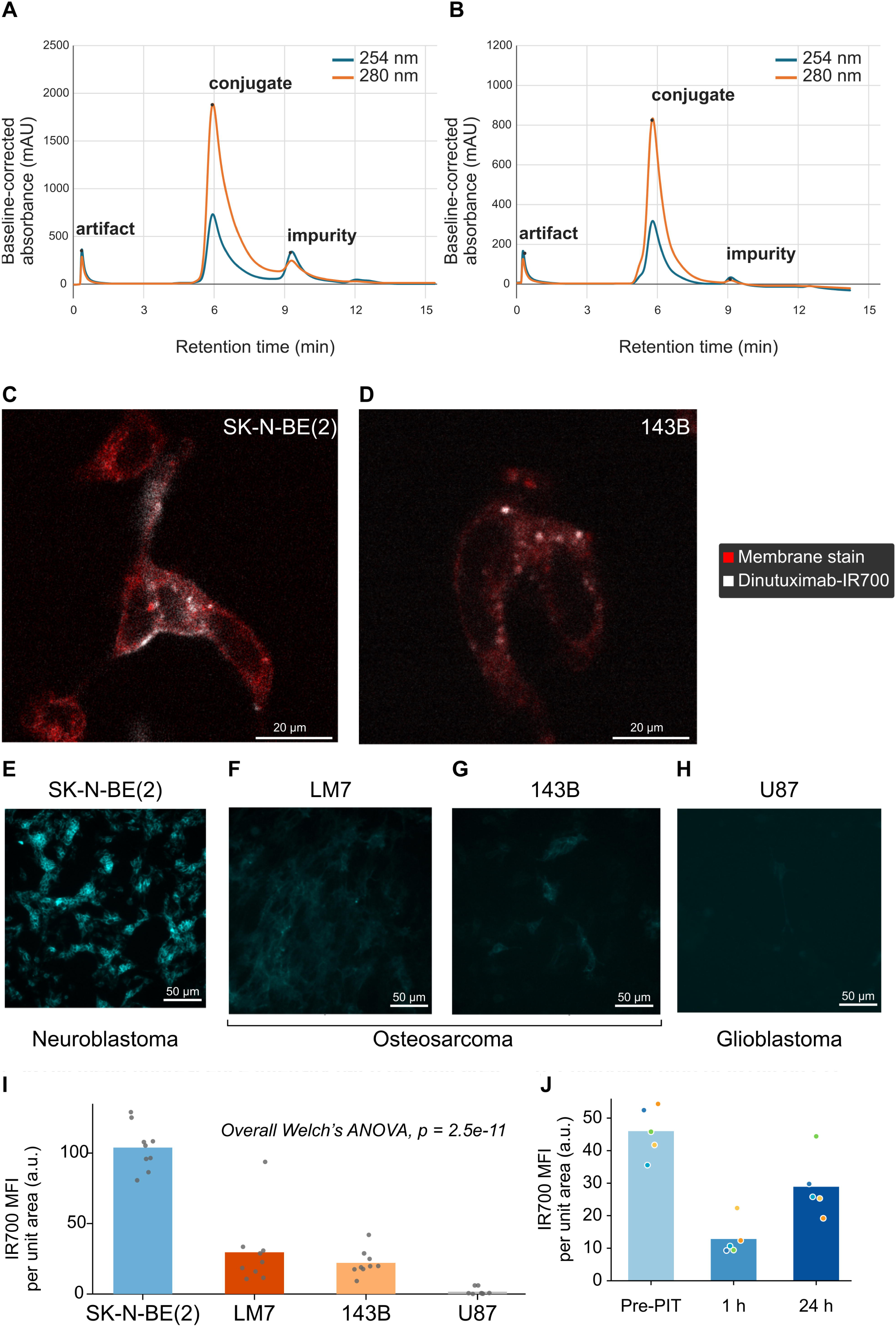
Chemical and fluorescence properties of dinutuximab-IR700. (A, B) SEC-HPLC characterization of dinutuximab-IR700 and dinutuximab-IR800 conjugates, respectively. (C, D) Confocal microscopy demonstrating dinutuximab-IR700 binding to NB and OS cell membranes, respectively. (E-H) IR700 fluorescence microscopy in NB, OS and GBM (I) IR700 MFI per unit area in NB, OS and GBM (J) Photobleaching of IR700 on PIT in SK-N-BE(2).

Confocal microscopy, performed to study tracer localization on GD2-expressing cells, demonstrated colocalization of dinutuximab-IR700 fluorescence with CellMask-defined plasma membranes in SK-N-BE(2) and 143B cells (Fig. 2C and D). Dinutuximab-IR700 bound more uniformly and consistently to SK-N-BE(2) cells than 143B cells, which also had subjectively sparse localization on the cell membrane.

Widefield fluorescence microscopy using the Cy5 channel enabled a comparison of the cell-associated IR700 signal in SK-N-BE(2), LM7, 143B, and U-87 MG cultures (Fig. 2E-I). Mean IR700 fluorescence intensity per unit area was highest in NB, intermediate in OS, and lowest in GBM (Welch’s one-way ANOVA, P = 2.5 × 10 ¹¹).

### PIT and PDT produced agent- and light-dose dependent killing

#### Dinutuximab-IR700 PIT

At a fixed dinutuximab-IR700 concentration of 15 µg/mL, increasing 689-nm light dose reduced viability in all four NB and OS cell lines (Fig. 3A-E). At 50 J/cm², mean viability was 23.6% ± 5.9%, 37.0% ± 5.9%, 47.3% ± 8.0%, and 79.6% ± 9.3% of control in NMB6, SK-N-BE(2), LM7, and 143B, respectively. Light dose significantly affected viability in NMB6, SK-N-BE(2), and LM7 (P_adj_ < 0.0001 for each). The response in 143B was less pronounced and more variable, although still significant (P_adj_ = 0.014). However, no individual light-dose versus 0 J/cm^2^ contrast remained significant after within-line Holm adjustment in 143B.

**Figure 3.**
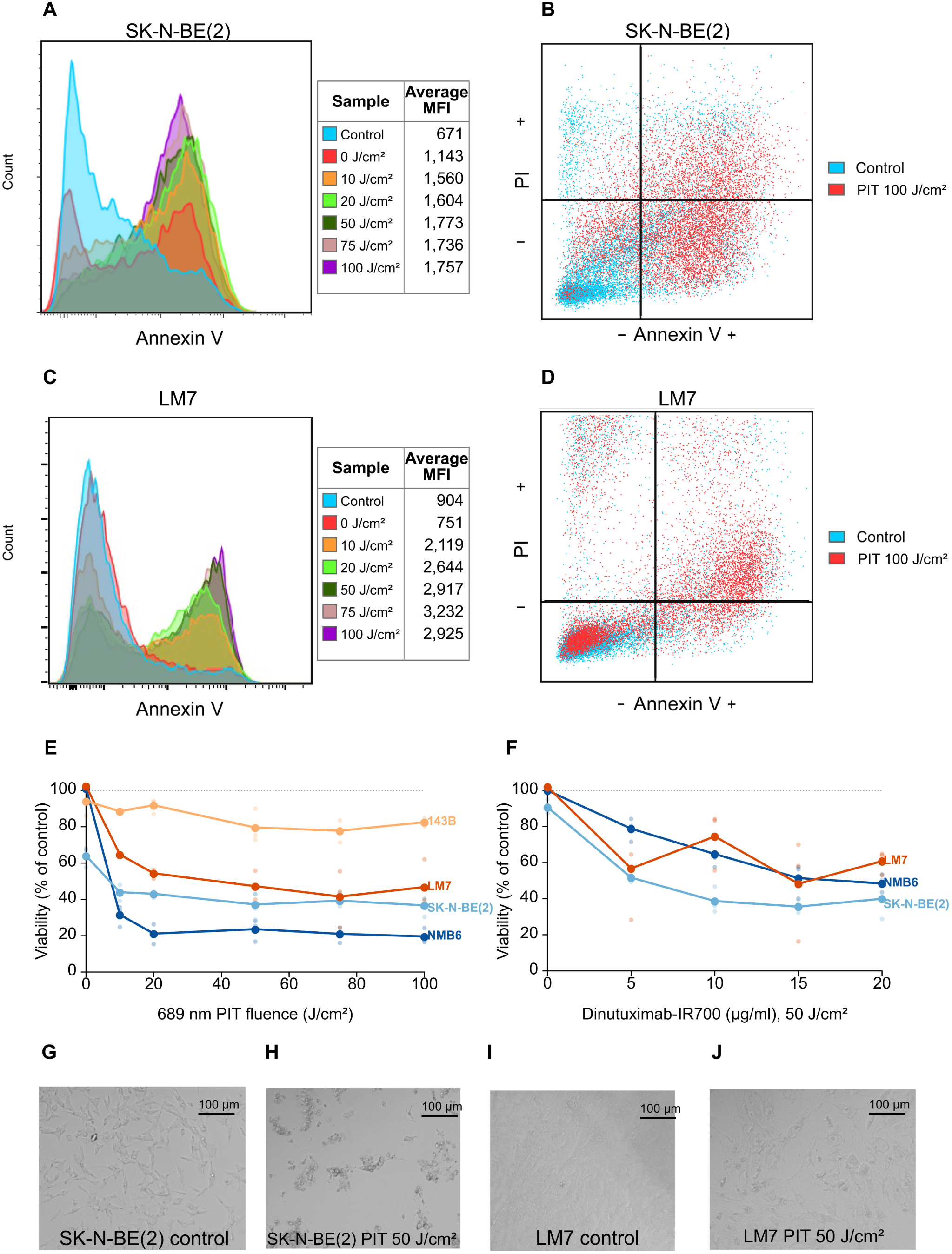
PIT monotherapy mediated tumor-cell killing. (A, C) Annexin V fluorescence histograms across PIT light doses in SK-N-BE(2) and LM7, respectively. (B, D) Representative Annexin V/propidium iodide plots comparing control cells with PIT at 100 J/cm² in SK-N-BE(2) and LM7, respectively. (E) Viability across 689-nm light doses at a fixed dinutuximab-IR700 concentration of 15 µg/mL. (F) Viability across dinutuximab-IR700 concentrations at a fixed 689-nm light dose of 50 J/cm². (G-J) Brightfield microscopy demonstrating cellular shrinkage and detachment in NB and OS, respectively, in response to PIT 50 J/cm^2^, compared with untreated control.

At a fixed light dose of 50 J/cm², increasing dinutuximab-IR700 concentration reduced viability in NMB6 and SK-N-BE(2) (P_adj_ < 0.0001 for each; Fig. 3F, S1A-D). Across 0-20 µg/mL, mean viability decreased from 100.0% to 48.4% of control in NMB6 and from 90.7% to 40.0% in SK-N-BE(2). LM7 also showed a significant overall concentration effect when tested through 30 µg/mL (P_adj_ < 0.0001), although the response was variable and nonmonotonic. Brightfield microscopy showed loss of adherent cellular architecture and accumulation of rounded or detached cells after PIT in SK-N-BE(2) and LM7 (Fig. 3G-J). At 100 J/cm² with 15 µg/mL dinutuximab-IR700, Annexin V-positive and Annexin V-PI double-positive events predominated in SK-N-BE(2) and LM7 plots, respectively (Fig. 3B and D).

#### 5-ALA PDT

Brightfield microscopy and Annexin V/PI flow cytometry showed treatment-associated cytotoxicity on PDT (Fig. 4A-D and G-J). At 500 µmol/L 5-ALA, increasing 635-nm light dose reduced viability in all four parental cell lines (Fig. 4E). At 30 J/cm², mean viability was 32.4% ± 4.6%, 46.2% ± 3.2%, 12.1% ± 3.9%, and 15.0% ± 11.3% of control in NMB6, SK-N-BE(2), LM7, and 143B, respectively. Similar to PIT, PDT light dose significantly affected viability across the tested doses in all four NB and OS cell lines (P_adj_ < 0.0001 for each).

**Figure 4.**
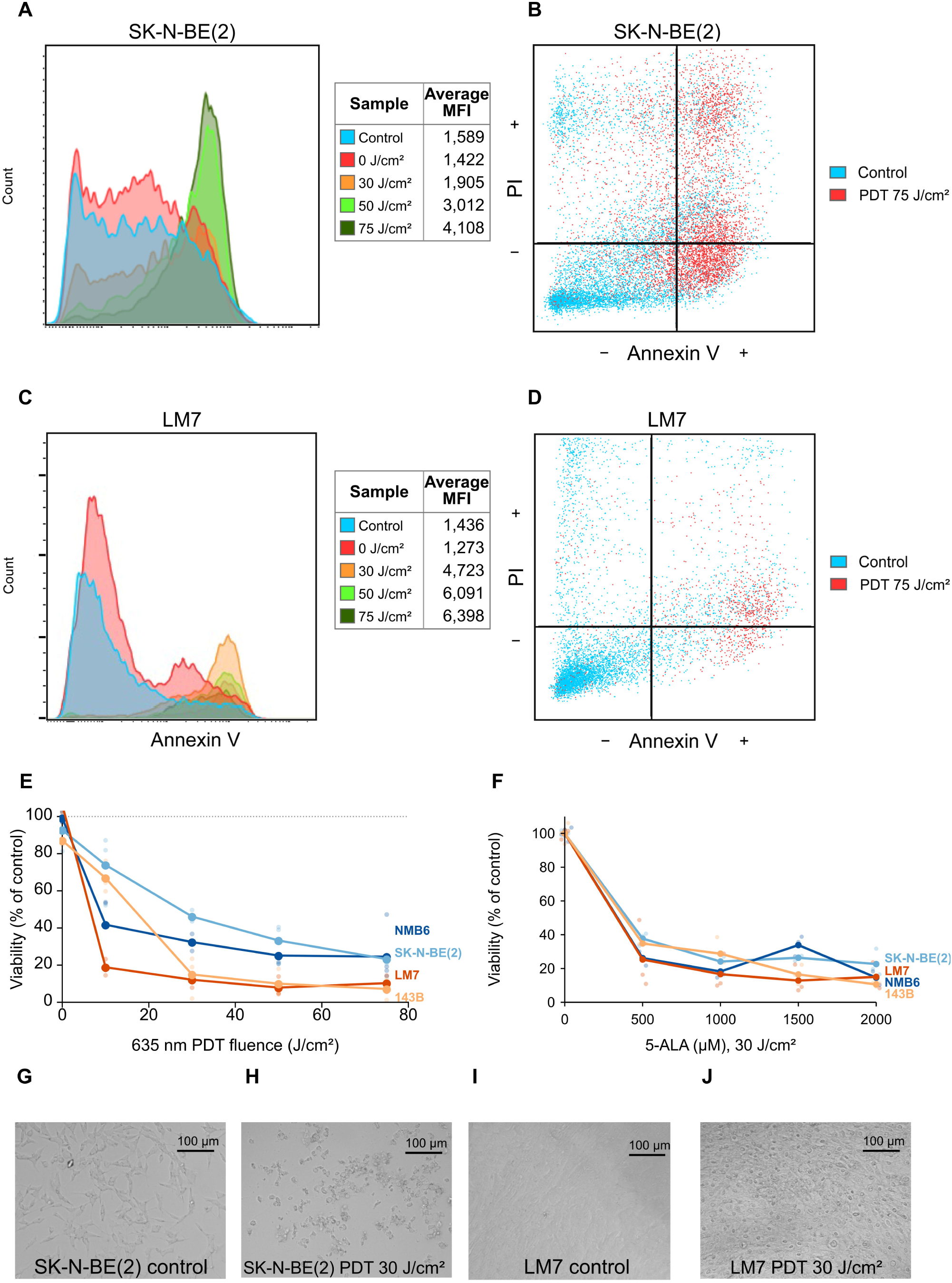
PDT monotherapy mediated tumor-cell killing. (A, C) Annexin V fluorescence histograms across PDT light doses in SK-N-BE(2) and LM7, respectively. (B, D) Representative Annexin V/propidium iodide plots comparing control cells with PDT at 75 J/cm² in SK-N-BE(2) and LM7, respectively. (E) Viability across 635-nm light doses at a fixed 5-ALA concentration of 500 µmol/L. (F) Viability across 5-ALA concentrations at a fixed 635-nm light dose of 30 J/cm². (G-J) Brightfield microscopy demonstrating cell shrinkage and detachment in NB and OS, respectively, on response to PDT 30 J/cm^2^, compared with untreated control.

The effect of increasing 5-ALA concentration at a fixed light dose of 30 J/cm² differed among cell lines (Fig. 4F, S1E-H). The within-cell-line concentration effect was significant in 143B (P_adj_ < 0.0001), in which mean viability decreased from 34.8% ± 0.4% of control at 500 µmol/L to 10.6% ± 1.1% at 2,000 µmol/L. The concentration effect did not reach significance in NMB6, SK-N-BE(2), or LM7 after adjustment across the four cell-line-specific tests (P_adj_ = 0.059, 0.059, and 0.874, respectively). Within each tested 5-ALA concentration, illuminated wells had lower viability than the corresponding 0 J/cm^2^ wells.

At the lowest light doses common to both light dose-response experiments, mean killing was greater with PIT than PDT in the NB cell lines, whereas PDT produced greater mean killing in OS.

### GD2 expression influences susceptibility to PIT

Owing to the antigen-specificity of PIT, a GD2 quantification study was undertaken for the PIT-treated cell lines. All four parental NB and OS cell lines expressed cell-surface GD2, although abundance differed among lines (Fig. 5A; Supplementary Table S2). Mean QuantiBRITE-derived estimates were 547,480 ± 23,663, 374,268 ± 16,709, 357,778 ± 11,647, and 154,413 ± 8,295 GD2 antigens per cell for NMB6, LM7, 143B, and SK-N-BE(2), respectively. SF8628 and U-87 MG, although GD2-expressing, had relatively lower estimates of 46,338 ± 1,402 and 17,136 ± 384, respectively.

**Figure 5.**
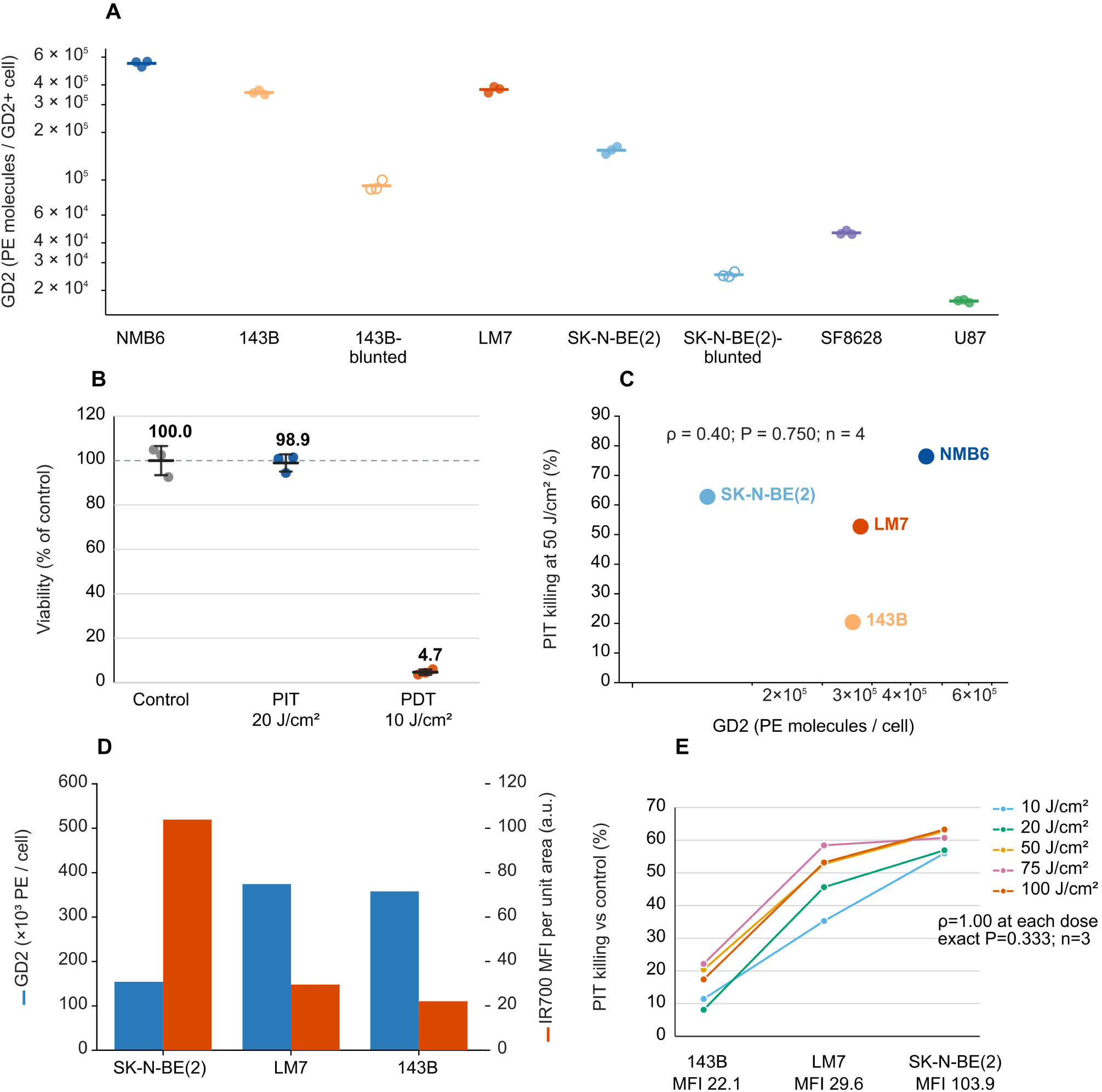
GD2 expression and cell-associated IR700 fluorescence in relation to PIT response (A) GD2 surface abundance across cell lines (B) Viability of HEK293T cells after PIT at 20 J/cm² or PDT at 10 J/cm² relative to untreated control (C) Control normalized PIT killing at 50 J/cm² in relation to GD2 expression in NB and OS (D) Per cell GD2 vs IR700 MFI per unit-area in SK-N-BE(2), LM7, and 143B (E) Rank order of IR700 MFI per unit area and PIT killing.

HEK293T, a cell line with no known GD2 expression, served as a control (Fig. 5B). Mean viability was 100.0% in untreated cells, 98.9% after dinutuximab-IR700 PIT at 20 J/cm², and 4.7% after 5-ALA PDT at 10 J/cm². PIT produced no measurable cytotoxicity, whereas PDT remained strongly cytotoxic. We further investigated the association between GD2 abundance and PIT killing in NB and OS at a fixed light dose of 50 J/cm² and dinutuximab-IR700 dose of 15 µg/mL. While a positive rank-order association was found, the relationship was modest and not statistically significant (Spearman ρ = 0.40; P =; n = 4; Fig. 5C).

On comparison of parental SK-N-BE(2) and 143B with their GD2-blunted derivatives, GD2 abundance was found to be reduced 6.1-fold in SK-N-BE(2) to 25,156 ± 948 GD2 antigens per cell, and 3.9-fold in 143B to 91,970 ± 7,099 (P_adj_ < 0.0001 for both; Fig. 5A). Notably, PIT did not reduce viability in either GD2-blunted derivative (SK-N-BE(2), P = 1.000; 143B, P = 1.000). However, PDT remained effective, reducing mean viability to 56.2% and 24.2% of control, respectively (P_adj_ < 0.0001; Supplementary Fig. S1K). These findings support GD2-dependent activity of dinutuximab-IR700 PIT and GD2-independence of PDT.

GD2 expression and IR700 fluorescence per image area followed different patterns (Fig. 5D). SK-N-BE(2) had the lowest calibrated GD2 expression among the three cell lines in the paired microscopy analysis but the highest IR700 fluorescence per image area. LM7 and 143B had higher per-cell GD2 estimates but lower fluorescence. While GD2 expression did not reproduce the observed PIT-response ordering, IR700 fluorescence precisely matched the rank order of PIT killing at every light dose (Spearman ρ = 1.00 at each dose; exact two-sided P = 0.333; n = 3; limited number of cell lines precluding statistical significance; Fig. 5E).

### Culture medium and time-dependence of PIT and PDT

In SK-N-BE(2) cells, absence of culture medium during illumination influenced the cytotoxicity mediated by the phototherapies. Removal of culture medium prior to illumination had opposite effects on the two phototherapies (P = 0.0010; Supplementary Fig. S2A-B). After PIT, mean viability was 42.1% ± 6.5% of control with medium and 87.6% ± 14.8% without medium (P_adj_ = 0.0080). PDT remained active without medium: mean viability was 44.5% ± 4.4% with medium and 27.0% ± 5.4% without medium (P_adj_ = 0.0080). Thus, transient removal of extracellular medium markedly attenuated PIT-mediated cytotoxicity but did not impair PDT under the tested conditions.

The temporal response also differed between modalities (P = 0.0035; Supplementary Fig. S2C-D). PIT reduced viability to 64.6% ± 4.9% of the matched control by 1 hour and to 52.0% ± 1.7% at 24 hours (1 vs. 24 hours, P_adj_ = 0.0099). Viability at 48 hours did not differ from the 24-hour value (P_adj_ = 0.099). In contrast, PDT viability progressively decreased from 83.5% ± 15.6% at 1 hour to 32.3% ± 5.4% at 24 hours and 22.0% ± 0.8% at 48 hours (P_adj_ = 0.0026 and 0.043, respectively). These data demonstrate an early PIT effect and a time-dependent increase in PDT cytotoxicity.

### Differential phototoxicity of IR700 and IR800 ADC

IR700 and IR800 were compared as photoabsorbers to confirm PIT toxicity rather than dinutuximab toxicity. Dinutuximab-IR700 reduced SK-N-BE(2) viability after 689-nm illumination, whereas dinutuximab-IR800 followed by 810-nm illumination produced little change from control (Supplementary Fig. S1I-J). At 24 hours, mean viability after IR700 treatment was approximately 53% of control at 20 J/cm² and 21%-23% at 50-100 J/cm², compared with approximately 90%-92% across the corresponding IR800 conditions.

Fluorescence-imaging studies further characterized factors associated with measured IR700 signal (Fig. 2J, Supplementary Fig. S1L-N, and Supplementary Fig. S2E-G). Mean fluorescence decreased from 46.0 arbitrary units before PIT to 12.8 at 1 hour and partially recovered to 28.9 at 24 hours, consistent with photobleaching. Imaging in culture medium yielded greater fluorescence than imaging without medium (59.9 vs. 20.8 units). Conjugates with 3 IR700 per antibody produced greater fluorescence than those with 2 (83.5 vs. 42.9 units), and fluorescence was greater with 20 µg than with 5 µg dinutuximab-IR700 (48.9 vs. 21.7 units).

### Cytotoxic effects and light-dose requirements of combined PIT and PDT

At 10 µg/mL dinutuximab-IR700 and 500 µmol/L 5-ALA, the global PIT-by-PDT interaction was significant in SK-N-BE(2) (P_adj_ < 0.0001) but not in LM7 (P_adj_ = 0.467). The significant interaction in SK-N-BE(2) indicates that the effect of one light modality changed across levels of the other within the dual-agent matrix (Fig. 6A and D). It, however, does not independently establish synergy. The sensitivity analysis supported the primary interaction findings, with significant PIT-by-PDT interaction in SK-N-BE(2) (P<0.0001) but not in LM7 (P =0.435).

**Figure 6.**
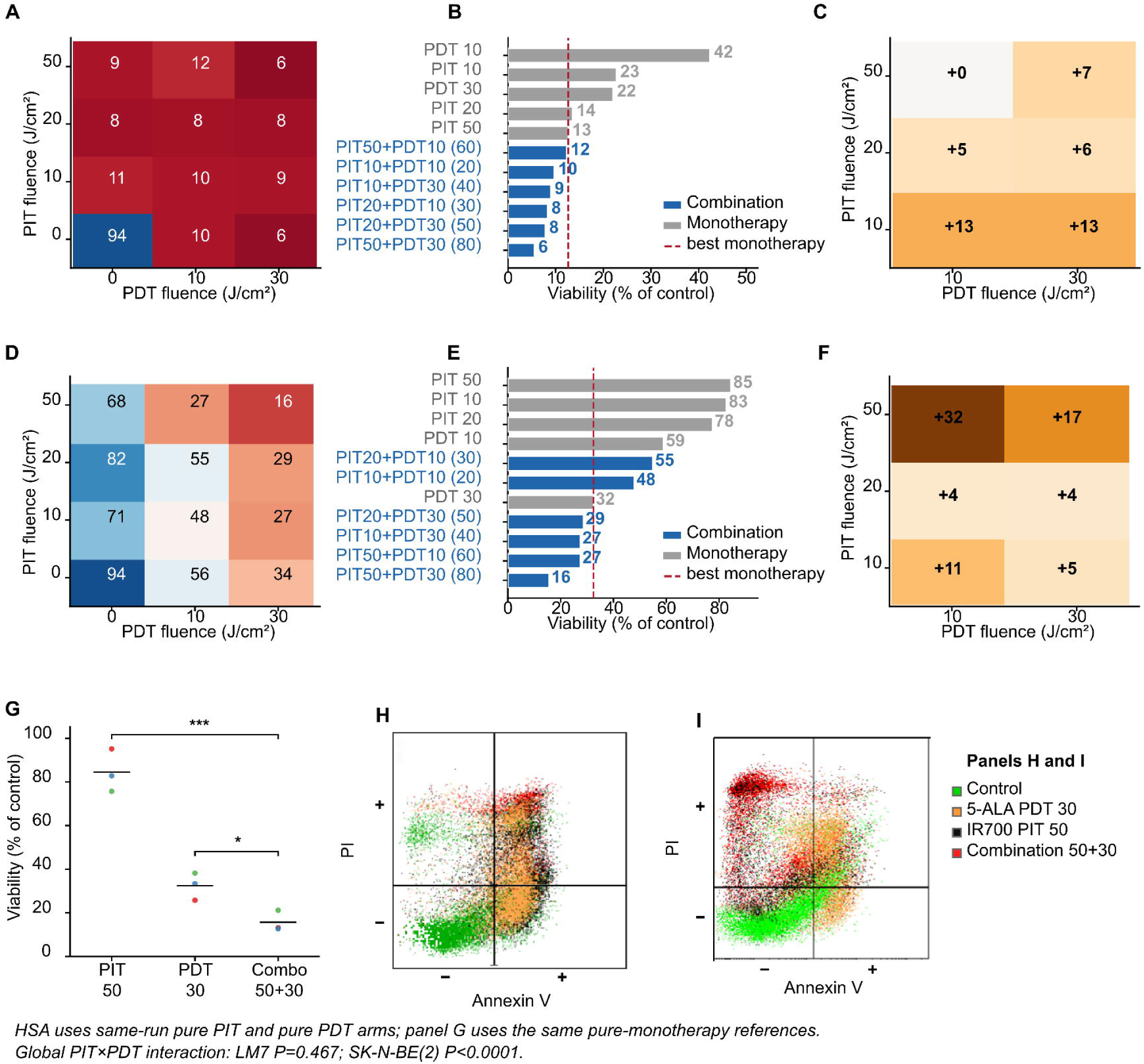
Combination PIT plus PDT at selected light-dose combinations. All light-doses are expressed in J/cm^2^. (A) SK-N-BE(2) viability on combination therapy (B) Light dose sparing effects of combination therapy in SK-N-BE(2) (C) SK-N-BE(2) HSA excess killing (percentage points) (D) LM7 viability on combination therapy (E) Light dose sparing effect of combination therapy in LM7 (F) LM7 HSA excess killing (percentage points) (G) LM7 combination vs pure monotherapy (H, I) Superimposed Annexin V/propidium iodide flow-cytometry events after control, 5-ALA PDT at 30 J/cm², dinutuximab-IR700 PIT at 50 J/cm², or combined treatment in.

Combined therapy improved cytotoxicity over pure monotherapy at selected light combinations. In SK-N-BE(2), PIT 10 J/cm² plus PDT 30 J/cm² and PIT 50 J/cm² plus PDT 30 J/cm² produced 13.1 and 7.1 percentage points of HSA excess killing, respectively, over corresponding monotherapies (P_adj_ = 0.0003 and 0.0022; Fig. 6C). In LM7, PIT 50 J/cm² plus PDT 30 J/cm² produced 16.8 percentage points of HSA excess killing (P_adj_ = 0.022; Fig. 6E-G). At this common regimen, mean viability was 5.6% ± 0.9% of control in SK-N-BE(2) and 15.7% ± 4.8% in LM7 (Supplementary Table S3).

Combined treatment also reduced the required light dose in SK-N-BE(2). All six dual-light combinations reduced mean viability to 5.6%-12.3% of control (Fig. 6A). PIT 10 J/cm² plus PDT 10 J/cm² produced 90.3% killing at a summed light dose of 20 J/cm², compared with 78.0% killing after PDT 30 J/cm² monotherapy and 87.0% killing with PIT 50 J/cm² monotherapy (Fig. 6B). At this lowest dual-light regimen, observed viability closely matched the Bliss-expected viability in SK-N-BE(2) (9.7% versus 9.6%; Bliss excess killing, −0.1 percentage points) and LM7 (47.9% versus 48.8%; +1.0 percentage point), consistent with Bliss additivity. Across the full matrices, Bliss excess killing ranged from −7.0 to −0.1 percentage points in SK-N-BE(2) and −9.2 to +22.4 percentage points in LM7, indicating dose- and cell-line-dependent interaction rather than uniform synergy. A factorial test evaluated whether the effect of either modality changed across the treatment matrix. SK-N-BE(2) showed a significant global interaction, while LM7 showed a nonsignificant global interaction. Together, these analyses support dual therapy in both cell lines but do not demonstrate uniform synergy.

Annexin V/PI flow cytometry showed depletion of the viable double-negative population after combined treatment in LM7 and SK-N-BE(2), accompanied by cell-line-specific increases in Annexin V-positive and/or PI-positive populations (Fig. 6H-I).

## DISCUSSION

Treatment of NB and OS presents notable surgical challenges (2, 9). Adequate resection of NB is often constrained by the close relationship of tumor to critical anatomical structures and difficulty distinguishing tumor from surrounding tissue (2). For OS, durable local control remains dependent on complete resection with negative margins, limited by involvement of crucial nerves, vessels and joints (9). Tumor heterogeneity further complicates treatment (31, 32). A therapeutic gap therefore remains for residual or incompletely resectable disease despite multimodal therapy (1, 6, 7, 9). Light-activated therapies are well suited to bridge this gap, through selective photosensitizer localization and targeted illumination (12, 15, 23, 24).

The National Cancer Institute has identified GD2 as one of the most crucial antigens for tumor-targeted therapies (33). GD2 is overexpressed on NB and OS with limited expression in normal tissues, but its abundance varies, and GD2-low states can confer dinutuximab resistance (18, 19, 34). We therefore paired GD2-directed PIT with antigen-independent 5-ALA PDT to extend cytotoxic coverage across heterogeneous tumor-cell populations. Conjugation and binding studies confirmed IR700 coupling and plasma-membrane localization. PIT was active across NB and OS cell lines, while limited cytotoxicity with dinutuximab-IR800 or without illumination supported dependence on the IR700-689-nm light interaction (12–15). This extends prior GD2-targeted PIT (20) using the clinically established dinutuximab backbone (6, 7).

Target dependence was demonstrated in GD2-blunted derivatives and HEK293T cells, in which PIT did not reduce viability whereas PDT remained effective. This supported a rationale for combination treatment. Among parental lines, GD2 abundance did not determine response. OS cells appeared larger on brightfield and confocal microscopy, suggesting that their GD2 abundance was distributed over a larger membrane area, likely yielding lower antigen and IR700 density and therefore lower cell-associated IR700 fluorescence per unit area. This may also explain why PIT killing followed IR700 fluorescence per unit area, since local density of membrane-bound APCs may influence membrane injury (13–15).

5-ALA PDT monotherapy was active in all four parental lines, consistent with previous NB and bone sarcoma studies (25–27). The decrease in HEK293T viability demonstrates potential PDT toxicity in non-tumor tissue. The medium and time-course experiments further distinguished PIT and PDT: PIT was markedly attenuated without culture medium, whereas PDT remained active. PIT reduced viability by 1 hour and further by 24 hours, while PDT cytotoxicity continued to increase through 48 hours. These differences in target dependence, treatment conditions, and kinetics support the biological rationale for combining PIT and PDT.

The combination experiments addressed whether these differences could improve treatment effect or reduce the required light exposure. In both SK-N-BE(2) and LM7, dual treatment exceeded matched monotherapy at selected doses. Moreover, in SK-N-BE(2), combination treatment with lower light dose (10 J/cm^2^) reduced tumor cell viability more than that observed with higher-light-dose monotherapy and was consistent with Bliss additivity. The same dose pair was also additive in LM7. The factorial interaction was significant in SK-N-BE(2) but not LM7, and Bliss excess killing was not consistently positive. Thus, the findings support a dose-specific combination benefit rather than uniform synergy. Further, substantial cytotoxicity can be achieved at lower summed light dose, shortening illumination time and surgical duration and potentially limiting phototherapy-mediated thermal injury (30) of surrounding normal tissue/organs.

Overall, this combined PIT and PDT approach produced substantial *in vitro* tumor-cell cytotoxicity, providing a strong basis for *in vivo* evaluation. Combination phototherapy holds promise as a potential surgical adjunct to ablate residual disease. This approach may be useful for exposed resection beds, close or anatomically constrained margins, and residual tumor that cannot be removed without substantial morbidity.

Limitations include the limited cohort of GD2-expressing tumors tested, unmeasured PpIX, ROS, tissue heating, and photosensitizer distribution, and use of immortalized HEK293T cell line rather than a normal-tissue model. Further three-dimensional, orthotopic, and patient-derived models should next examine delivery, light penetration, normal-tissue effects and immune responses associated with phototherapy. Although the exclusively *in vitro* design is a limitation, validation across multiple experimental conditions *in vitro* provides a foundation for future *in vivo* testing.

While there have been advances in phototherapy, its use in pediatric extracranial tumors remains limited. We demonstrated the *in vitro* efficacy of a dual-phototherapy combination regimen, which enhanced cytotoxicity relative to either monotherapy at selected doses while maintaining substantial tumor-cell killing at lower summed light dose. These findings support further *in vivo* evaluation of PIT, PDT, and combination therapy as a spatially controlled local treatment for residual tumors with heterogeneous GD2 expression and treatment sensitivity.

## ACKNOWLEDGMENTS

The authors acknowledge the Flow Cytometry Core at John G. Rangos Sr. Research Center, the University of Pittsburgh Dietrich School Microscopy and Imaging Suite and the UPMC Hillman Cancer Center Small Animal Multimodal Imaging Facility for technical support. Generative AI was used to aid with figure design.

## AUTHOR CONTRIBUTIONS

AP contributed to the conceptualization, data curation, formal analysis, investigation, methodology, project administration, software, validation, visualization, writing, review and editing of this paper. CY and AK contributed to the review and editing of this paper. MM contributed to the resources, software and investigation. BC, VK, EN, BG and SJ contributed to the investigation. PC and HK contributed to the resources, review and editing. CH contributed to the resources, software, review and editing. LR contributed to the conceptualization, investigation, methodology, review and editing. MMM and GK contributed to the conceptualization, funding acquisition, investigation, methodology, project administration, resources, software, supervision, review and editing of this paper.

---

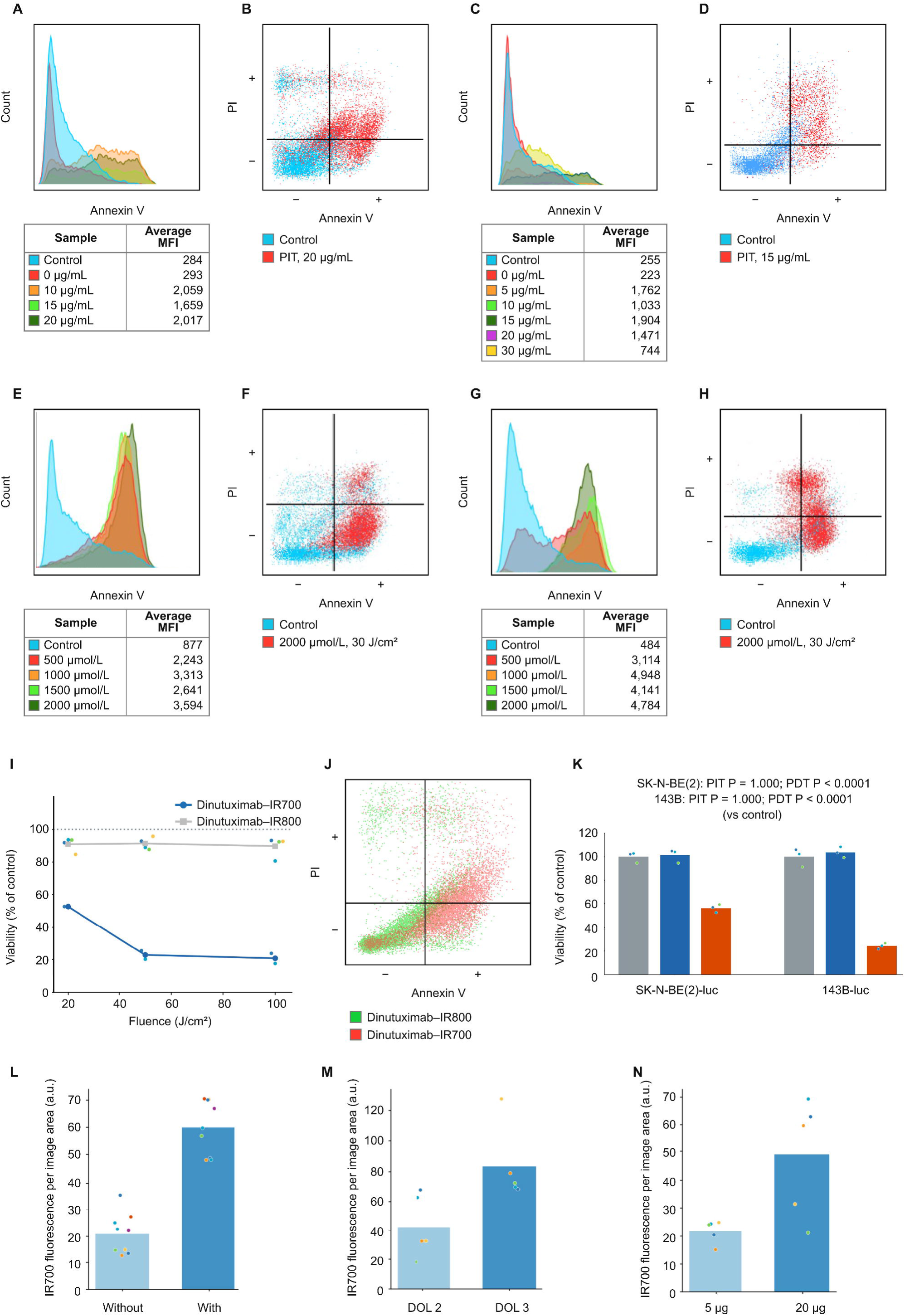

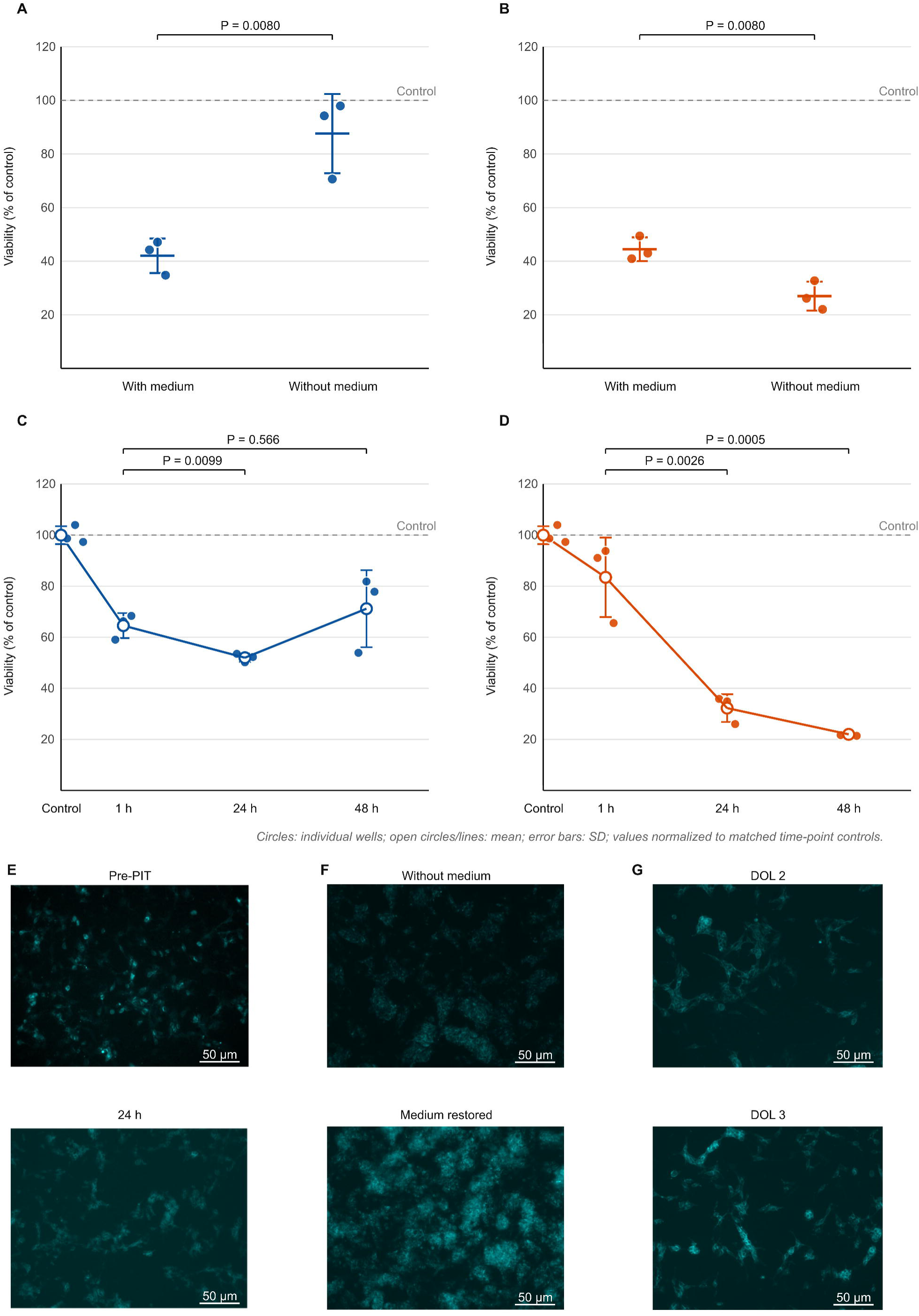

## REFERENCES

1. Maris JM, Hogarty MD, Bagatell R, Cohn SL. Neuroblastoma. Lancet 2007;369:2106–20.

2. Rosenblum LT, Sever RE, Gilbert R, Guerrero D, Vincze SR, Menendez DM, et al. Dual-labeled anti-GD2 targeted probe for intraoperative molecular imaging of neuroblastoma. J Transl Med 2024;22:940.

3. Temple WC, Vo KT, Matthay KK, Balliu B, Coleman C, Michlitsch J, et al. Association of image-defined risk factors with clinical features, histopathology, and outcomes in neuroblastoma. Cancer Med 2021;10:2232–41.

4. Espinoza AF, Bagatell R, McHugh K, Naranjo AH, Van Ryn C, Rojas Y, et al. A subset of image-defined risk factors predict completeness of resection in children with high-risk neuroblastoma: an international multicenter study. Pediatr Blood Cancer 2024;71:e31218.

5. von Allmen D, Davidoff AM, London WB, Van Ryn C, Haas-Kogan DA, Kreissman SG, et al. Impact of extent of resection on local control and survival in patients from the COG A3973 study with high-risk neuroblastoma. J Clin Oncol 2017;35:208–16.

6. Yu AL, Gilman AL, Ozkaynak MF, London WB, Kreissman SG, Chen HX, et al. Anti-GD2 antibody with GM-CSF, interleukin-2, and isotretinoin for neuroblastoma. N Engl J Med 2010;363:1324–34.

7. Yu AL, Gilman AL, Ozkaynak MF, Naranjo A, Diccianni MB, Gan J, et al. Long-term follow-up of a phase III study of ch14.18 (dinutuximab) + cytokine immunotherapy in children with high-risk neuroblastoma: COG study ANBL0032. Clin Cancer Res 2021;27:2179–89.

8. London WB, Castel V, Monclair T, Ambros PF, Pearson ADJ, Cohn SL, et al. Clinical and biologic features predictive of survival after relapse of neuroblastoma: a report from the International Neuroblastoma Risk Group project. J Clin Oncol 2011;29:3286–92.

9. Isakoff MS, Bielack SS, Meltzer P, Gorlick R. Osteosarcoma: current treatment and a collaborative pathway to success. J Clin Oncol 2015;33:3029–35.

10. Pastorino U, Palmerini E, Porcu L, Luksch R, Scanagatta P, Meazza C, et al. Lung metastasectomy for osteosarcoma in children, adolescents, and young adults: proof of permanent cure. Tumori 2023;109:79–85.

11. Spraker-Perlman HL, Barkauskas DA, Krailo MD, Meyers PA, Schwartz CL, Doski J, et al. Factors influencing survival after recurrence in osteosarcoma: a report from the Children’s Oncology Group. Pediatr Blood Cancer 2019;66:e27444.

12. Mitsunaga M, Ogawa M, Kosaka N, Rosenblum LT, Choyke PL, Kobayashi H. Cancer cell-selective in vivo near infrared photoimmunotherapy targeting specific membrane molecules. Nat Med 2011;17:1685–91.

13. Sato K, Ando K, Okuyama S, Moriguchi S, Ogura T, Totoki S, et al. Photoinduced ligand release from a silicon phthalocyanine dye conjugated with monoclonal antibodies: a mechanism of cancer cell cytotoxicity after near-infrared photoimmunotherapy. ACS Cent Sci 2018;4:1559–69.

14. Nakajima K, Takakura H, Shimizu Y, Ogawa M. Changes in plasma membrane damage inducing cell death after treatment with near-infrared photoimmunotherapy. Cancer Sci 2018;109:2889–96.

15. Kobayashi H, Griffiths GL, Choyke PL. Near-infrared photoimmunotherapy: photoactivatable antibody-drug conjugates (ADCs). Bioconjug Chem 2020;31:28–36.

16. Cognetti DM, Johnson JM, Curry JM, Kochuparambil ST, McDonald D, Mott F, et al. Phase 1/2a, open-label, multicenter study of RM-1929 photoimmunotherapy in patients with locoregional, recurrent head and neck squamous cell carcinoma. Head Neck 2021;43:3875–87.

17. Gomes-da-Silva LC, Kepp O, Kroemer G. Regulatory approval of photoimmunotherapy: photodynamic therapy that induces immunogenic cell death. Oncoimmunology 2020;9:1841393.

18. Roth M, Linkowski M, Tarim J, Piperdi S, Sowers R, Geller D, et al. Ganglioside GD2 as a therapeutic target for antibody-mediated therapy in patients with osteosarcoma. Cancer 2014;120:548–54.

19. Anderson J, Majzner RG, Sondel PM. Immunotherapy of neuroblastoma: facts and hopes. Clin Cancer Res 2022;28:3196–206.

20. Inagaki FF, Kato T, Furusawa A, Okada R, Wakiyama H, Furumoto H, et al. Disialoganglioside GD2-targeted near-infrared photoimmunotherapy (NIR-PIT) in tumors of neuroectodermal origin. Pharmaceutics 2022;14:2037.

21. Zhao J, Nakajima K, Ueki M, Terashita Y, Hirabayashi S, Cho Y, et al. Near-infrared photoimmunotherapy using antiGD2 antibody for neuroblastoma and osteosarcoma. EJC Paediatr Oncol 2025;5:100233.

22. Nouso H, Tazawa H, Tanimoto T, Tani M, Watanabe H, Oyama T, et al. Near-infrared photoimmunotherapy targeting high-risk human neuroblastoma cells expressing GD2. Anticancer Res 2026;46:25–38.

23. Mroz P, Yaroslavsky A, Kharkwal GB, Hamblin MR. Cell death pathways in photodynamic therapy of cancer. Cancers (Basel) 2011;3:2516–39.

24. Casas A. Clinical uses of 5-aminolaevulinic acid in photodynamic treatment and photodetection of cancer: a review. Cancer Lett 2020;490:165–73.

25. Bergmann F, Stepp H, Metzger R, Rolle U, Johansson A, Till H. In vitro and in vivo evaluation of photodynamic techniques for the experimental treatment of human hepatoblastoma and neuroblastoma: preliminary results. Pediatr Surg Int 2008;24:1331–33.

26. Okamura SM, Chelakkot VS, Linn Z, Onishi Y, Nakahata K, Toyoda H, et al. 6-Aminonicotinamide enhances the efficacy of 5-aminolevulinic acid-mediated photodynamic therapy for neuroblastoma. BMC Cancer 2025;25:1815.

27. Maggs RH, Brookes MJ, Rankin KS. An in vitro investigation of 5-aminolevulinic acid mediated photodynamic therapy in bone sarcoma. Oncol Res 2026;34:1.

28. Vermandel M, Dupont C, Lecomte F, Leroy HA, Tuleasca C, Mordon S, et al. Standardized intraoperative 5-ALA photodynamic therapy for newly diagnosed glioblastoma patients: a preliminary analysis of the INDYGO clinical trial. J Neurooncol 2021;152:501–14.

29. Peciu-Florianu I, Vannod-Michel Q, Vauleon E, Bonneterre ME, Reyns N. Long term follow-up of patients with newly diagnosed glioblastoma treated by intraoperative photodynamic therapy: an update from the INDYGO trial (NCT03048240). J Neurooncol 2024;168:495–505.

30. Okuyama S, Nagaya T, Ogata F, Maruoka Y, Sato K, Nakamura Y, et al. Avoiding thermal injury during near-infrared photoimmunotherapy (NIR-PIT): the importance of NIR light power density. Oncotarget 2017;8:113194–201.

31. Schmelz K, Toedling J, Huska M, Cwikla MC, Kruetzfeldt LM, Proba J, et al. Spatial and temporal intratumour heterogeneity has potential consequences for single biopsy-based neuroblastoma treatment decisions. Nat Commun 2021;12:6804.

32. Kinnaman MD, Zaccaria S, Makohon-Moore A, Arnold B, Levine MF, Gundem G, et al. Subclonal somatic copy-number alterations emerge and dominate in recurrent osteosarcoma. Cancer Res 2023;83:3796–812.

33. Cheever MA, Allison JP, Ferris AS, Finn OJ, Hastings BM, Hecht TT, et al. The prioritization of cancer antigens: a National Cancer Institute pilot project for the acceleration of translational research. Clin Cancer Res 2009;15:5323–37.

34. Mabe NW, Huang M, Dalton GN, Alexe G, Schaefer DA, Geraghty AC, et al. Transition to a mesenchymal state in neuroblastoma confers resistance to anti-GD2 antibody via reduced expression of ST8SIA1. Nat Cancer 2022;3:976–93.

