## Supplementary tables and supplementary figure legends for "Complementary cytotoxicity of GD2-targeted photoimmunotherapy and 5-aminolevulinic acid photodynamic therapy in neuroblastoma and osteosarcoma"

### **SUPPLEMENTARY MATERIAL**

#### **Supplementary Table S1.** Experimental conditions for monotherapy dose-response studies

| **Modality** | **Dose-response experiment** | **Cell line(s)** | **Agent concentrations** | **Light exposure** | **Fixed condition** |
| --- | --- | --- | --- | --- | --- |
| PIT | Light-dose response | NMB6, SK-N-BE(2), LM7, and 143B | 15 µg/mL dinutuximab-IR700 | 0, 10, 20, 50, 75, and 100 J/cm² at 689 nm | Dinutuximab-IR700 fixed at 15 µg/mL |
| PIT | Agent-dose response | NMB6 and SK-N-BE(2) | 0, 5, 10, 15, and 20 µg/mL dinutuximab-IR700 | 50 J/cm² at 689 nm | Light dose fixed at 50 J/cm² |
| PIT | Agent-dose response | LM7 | 0, 5, 10, 15, 20, and 30 µg/mL dinutuximab-IR700 | 50 J/cm² at 689 nm | Light dose fixed at 50 J/cm² |
| PDT | Light-dose response | NMB6, SK-N-BE(2), LM7, and 143B | 500 µmol/L 5-ALA | 0, 10, 30, 50, and 75 J/cm² at 635 nm | 5-ALA fixed at 500 µmol/L |
| PDT | Agent-dose response | NMB6, SK-N-BE(2), LM7, and 143B | 0, 500, 1,000, 1,500, and 2,000 µmol/L 5-ALA | 0 or 30 J/cm² at 635 nm | Illumination evaluated at 0 and 30 J/cm² |

#### **Supplementary Table S2.** Cell-line panel and cell-surface GD2 abundance

| Cell line | Tumor type/role | Cell-line type | Antibody-associated PE molecules per GD2-positive cell, mean ± SD |
| --- | --- | --- | --- |
| NMB6 | NB | Parental | 547,480 ± 23,663 |
| SK-N-BE(2) | NB | Parental | 154,413 ± 8,295 |
| SK-N-BE(2), GD2-blunted | NB | GD2-blunted derivative | 25,156 ± 948 |
| LM7 | OS | Parental | 374,268 ± 16,709 |
| 143B | OS | Parental | 357,778 ± 11,647 |
| 143B, GD2-blunted | OS | GD2-blunted derivative | 91,970 ± 7,099 |
| SF8628 | Diffuse midline glioma | Parental | 46,338 ± 1,402 |
| U-87 MG | GBM | Parental | 17,136 ± 384 |

#### **Supplementary Table S3.** Same-run pure-monotherapy and combination viability at a common light regimen

| **Cell line** | **Regimen** | **PIT light dose, J/cm²** | **PDT light dose, J/cm²** | **Viability, % of control, mean ± SD** |
| --- | --- | --- | --- | --- |
| SK-N-BE(2) | PIT-only | 50 | 0 | 12.7 ± 2.6 |
| SK-N-BE(2) | PDT-only | 0 | 30 | 22.0 ± 3.2 |
| SK-N-BE(2) | PIT+PDT | 50 | 30 | 5.6 ± 0.9 |
| LM7 | PIT-only | 50 | 0 | 84.6 ± 9.8 |
| LM7 | PDT-only | 0 | 30 | 32.4 ± 6.3 |
| LM7 | PIT+PDT | 50 | 30 | 15.7 ± 4.8 |

### **SUPPLEMENTARY FIGURE LEGENDS**

**Figure S1.** Monotherapy flow-cytometry profiles and treatment-characterization substudies. (A, C) Annexin V fluorescence histograms across dinutuximab-IR700 concentrations in SK-N-BE(2) and LM7, respectively. (B, D) Representative Annexin V/propidium iodide plots comparing control with PIT at 20 µg/mL in SK-N-BE(2) and 15 µg/mL in LM7, respectively, using 50 J/cm² at 689 nm. (E, G) Annexin V fluorescence histograms across 5-ALA concentrations in SK-N-BE(2) and LM7, respectively. (F, H) Representative Annexin V/propidium iodide plots comparing control with PDT using 2,000 µmol/L 5-ALA and 30 J/cm² in SK-N-BE(2) and LM7, respectively. (I, J) Viability and representative flow-cytometry events after dinutuximab-IR700 with 689-nm illumination or dinutuximab-IR800 with 810-nm illumination in SK-N-BE(2). (K) Viability of GD2-blunted SK-N-BE(2) and 143B derivatives after PIT or PDT relative to control. (L-N) IR700 fluorescence per image area with and without culture medium, for conjugates with 2 and 3 IR700 per antibody, and after exposure to 5 or 20 µg dinutuximab-IR700, respectively.

**Figure S2.** Culture-medium dependence, temporal response, and representative fluorescence images. (A, B) Control-normalized viability after PIT and PDT, respectively, with or without culture medium during illumination. (C, D) Control-normalized viability at 1, 24, and 48 hours after PIT and PDT, respectively. (E) Representative widefield Cy5-channel images before PIT and 24 hours after illumination. (F) Representative Cy5-channel images obtained without culture medium and after medium restoration. (G) Representative Cy5-channel images obtained using dinutuximab-IR700 conjugates with 2 and 3 IR700 per antibody.
